# Extensive non-canonical phosphorylation in human cells revealed using strong-anion exchange-mediated phosphoproteomics

**DOI:** 10.1101/202820

**Authors:** Gemma Hardman, Simon Perkins, Zheng Ruan, Natarajan Kannan, Philip Brownridge, Dominic P Byrne, Patrick A. Eyers, Andrew R. Jones, Claire E. Eyers

**Author notes:** Corresponding author: Prof. Claire E. Eyers, Centre for Proteome Research, University of Liverpool.

## Abstract

Protein phosphorylation is a ubiquitous post-translational modification (PTM) that regulates all aspects of life. To date, investigation of human cell signalling has focussed on canonical phosphorylation of serine (Ser), threonine (Thr) and tyrosine (Tyr) residues. However, mounting evidence suggests that phosphorylation of histidine also plays a central role in regulating cell biology. Phosphoproteomics workflows rely on acidic conditions for phosphopeptide enrichment, which are incompatible with the analysis of acid-labile phosphorylation such as histidine. Consequently, the extent of non-canonical phosphorylation is likely to be under-estimated.

We report an Unbiased Phosphopeptide enrichment strategy based on Strong Anion Exchange (SAX) chromatography (UPAX), which permits enrichment of acid-labile phosphopeptides for characterisation by mass spectrometry. Using this approach, we identify extensive and positional phosphorylation patterns on histidine, arginine, lysine, aspartate and glutamate in human cell extracts, including 310 phosphohistidine and >1000 phospholysine sites of protein modification. Remarkably, the extent of phosphorylation on individual non-canonical residues vastly exceeds that of basal phosphotyrosine. Our study reveals the previously unappreciated diversity of protein phosphorylation in human cells, and opens up avenues for exploring roles of acid-labile phosphorylation in any proteome using mass spectrometry.

## Introduction

It is established that post-translational modification (PTM) of proteins by the addition of phosphate rapidly and reversibly regulates signalling functions, including catalytic activity, protein:protein interactions, protein localisation and stability^1, 2^. Consequently, phosphorylation is an essential PTM that is observed across the kingdoms of life, with dysregulation in humans being associated with diseases such as cancer. Until recently, phosphorylation of the hydroxyl-containing amino acids serine (Ser), threonine (Thr) and tyrosine (Tyr) was thought to be the primary mode of phosphorylation-mediated signalling in non-plant eukaryotes. However, a growing body of evidence^3-13^ indicates that phosphorylation on the imidazole-containing amino acid histidine (His) also plays a significant, but ill-defined, role in human biology. In contrast, and despite occasional reports in the literature^14, 15^, the prevalence (or otherwise) of non-canonical amino acid phosphorylation on the five amino acids for which protein phosphorylation is chemically feasible (Asp, Arg, Lys, Glu, Cys) has yet to be revealed in human cells.

Unlike the relatively stable phosphate esters (pSer, pThr, pTyr), the phosphoramidate bond in phosphohistidine (pHis) is susceptible to hydrolysis at non-physiological pH and/or elevated temperature^16-19^. Consequently, traditional (bio)chemical techniques typically used for characterising polypeptides containing phosphorylated Ser, Thr or Tyr, including denaturation with heat and acid-based MS workflows, are unsuitable for analysis of pHis and other ‘atypical’ phosphorylated amino acids, since the phosphate group is eliminated before the site of peptide modification can be defined^20^. A cellular understanding of the roles of acid-labile protein phosphorylation in vertebrates therefore lags far behind that of pSer, pThr and pTyr, and focussed analytical tools are essential to help redress this imbalance^6, 20-22^.

Recent development of generic antibodies recognising 1-pHis or 3-pHis isomers on proteins^12, 23, 24^, has revealed extensive cell-type independent modification^12^, and significant, but as yet incompletely defined roles for pHis have been proposed in processes as diverse as proliferation, differentiation and migration, ion channel regulation^13^, and T-cell signalling^7^. However, although pHis antibodies have been useful in facilitating the identification of novel pHis-containing proteins in human cells, specific sites of modification have not been defined, and these reagents are unsuitable for experimental analysis of other acid-labile protein phosphorylation events. To address these significant limitations, we developed an Unbiased Phosphopeptide enrichment strategy based on Strong Anion Exchange (SAX) chromatography (UPAX), which permits enrichment of acid-labile phosphopeptides at near neutral pH. This led to the facile identification of ^~^310 novel sites of cellular His phosphorylation from human cell extracts, and characterisation of pHis motifs in proteins. Moreover, UPAX permitted the identification and site localisation of hundreds of non-canonical protein phosphorylation events on the common amino acids Asp, Arg, Glu and Lys, which were previously assumed to lack extensive phosphorylation in vertebrate cells. Application and further development of the UPAX workflow will be of significant utility in understanding both canonical and non-canonical phosphorylation-mediated signalling from a global perspective, rather than the phosphoester-centric view that has traditionally dominated our understanding of human cell-based signalling.

## Results

### pH stability of phosphohistidine-containing peptides

Histidine phosphorylated peptides cannot be synthesised using standard solid-phase approaches. However, facile chemical phosphorylation of peptide and protein (*e.g.* myoglobin) histidine residues can be achieved using potassium phosphoramidate, generating suitable analytical standards^20, 25-28^. Intact protein analysis of phosphoramidate-treated myoglobin demonstrated up to five sites of His modification (Supp. methods; Supp. Fig. 1). Analysis of tryptic peptides revealed five pHis-containing peptides, which were mapped to seven sites of His phosphorylation in myoglobin (Supp. Fig. 2; Supp. Table 1). Although stability analysis has been reported for free pHis amino acids^18^, an assessment of pHis peptide stability over a range of pH values has not been undertaken. Phosphopeptide-enrichment prior to MS is traditionally performed at acidic pH, so we first assessed the pH-dependence of pHis peptide stability (Supp. Fig. 3). Myoglobin pHis tryptic peptides were much more tolerant of mildly acidic conditions than previously assumed, with a t_1/2_ at pH 4 of >2 hours, comparable to that observed at near neutral pH values. However, none of the pHis-peptides were stable at pH 1, with t_1/2_ values of ^~^15 min. Surprisingly, myoglobin-derived pHis peptides also exhibited decreased stability at pH 9, with a t_1/2_ of ^~^ 2 hours, potentially narrowing the pH window over which these phosphopeptides can be evaluated.

### Standard phosphopeptide enrichment strategies are unsuitable for pHis-containing peptides

Given the relative stability of pHis peptides at mildly acidic pH, particularly over short time periods, we compared pHis-peptide enrichment and recovery using standard isolation strategies. Tryptic peptides from His-phosphorylated myoglobin in a background of α-/β-casein-derived pSer/pThr peptides were subjected to rapid TiO_2_ enrichment under standard (low pH; Supp. Table 2, condition A), and non-standard (elevated pH, Supp. Table 2, conditions B, C) conditions. Condition C was based on a method described for analysis of the acid-labile amino acid phosphoarginine (pArg)^29^. Interestingly, none of the five pHis myoglobin peptides were observed following enrichment under any of the conditions assessed, even though phosphopeptide binding approached 100% (Supp. Table 3) and recovery of ≤95% was observed for pSer/Thr-containing phosphopeptides. Increased levels of non-phosphorylated peptides in the eluent compared to the phosphorylated starting material (*e.g*. Supp. Fig. 4), confirmed that peptide dephosphorylation during the enrichment protocol was the primary reason for the lack of pHis peptides identified.

We next evaluated calcium phosphate-based phosphopeptide enrichment, and hydroxyapatite (HAP) chromatography, both of which have been used for phosphopeptide isolation^30-32^. HAP was of particular interest as the pH is maintained at 7.2-7.8 throughout. However, for differing reasons the recovery of pHis-containing peptides using these two methods was very poor (see Supp. methods and results) and these purification procedures were deemed unsuited to pHis analysis.

### Strong anion exchange for enrichment of acid-labile pHis-containing peptides

Ion exchange chromatography, in which peptides are separated based on differences in charge, has been used extensively for peptide fractionation prior to LC-MS/MS-based proteomics^33^. Although strong cation exchange (SCX) is usually preferred for total proteome analyses^34-37^, strong anion exchange (SAX) is also suited to phosphopeptide separation, given the high abundance of negatively charged phosphate groups, including a second formal charge acquired above pH ^~^6^38, 39^. However, phosphopeptide elution from SAX columns is typically performed by a decrease in pH, potentially compromising pHis peptide stability. To assess the effect of ionic strength (as opposed to pH-mediated elution) for phospho(His)peptide analysis, we employed a modified triethylammonium phosphate gradient^38^, for phosphopeptide separation. As shown in Fig. 1A, the majority of phosphopeptide ion signals eluted in later SAX fractions, while non-phosphorylated peptides dominated earlier fractions (Fig. 1a). Fractionation of this phosphopeptide mixture by high pH reverse-phase chromatography confirmed that differential chromatographic elution of phosphopeptides from non-phosphopeptides is not simply an effect of peptide separation, but arises due to charge-based differences in affinity for the SAX solid phase. Extracted ion chromatograms (XICs) of the pHis peptides and their non-phosphorylated counterparts revealed that some of the pHis peptides underwent neutral loss over the time-course of the separation when SAX was performed at pH 6.0 (Supp. Fig. 5). Optimal separation of pHis peptides from their non-phosphorylated counterparts was readily observed at near-physiological pH (Fig. 1b, Supp. Fig. 5), with the latter SAX fractions being enriched for this simple phosphopeptide mixture by up to 100%. Importantly, spiking phosphorylated myoglobin into a complex human cell lysate did not affect separation of pHis peptides by SAX under these conditions. Moreover, LC-MS/MS analysis confirmed that recovery of pHis myoglobin peptides was similar to that of the non-phosphorylated and pSer/pThr peptides from of a ‘spiked’ standard mixture. Together, these data validate UPAX for phospho(His)peptide separation and analysis from complex mixtures.

**Figure 1.**
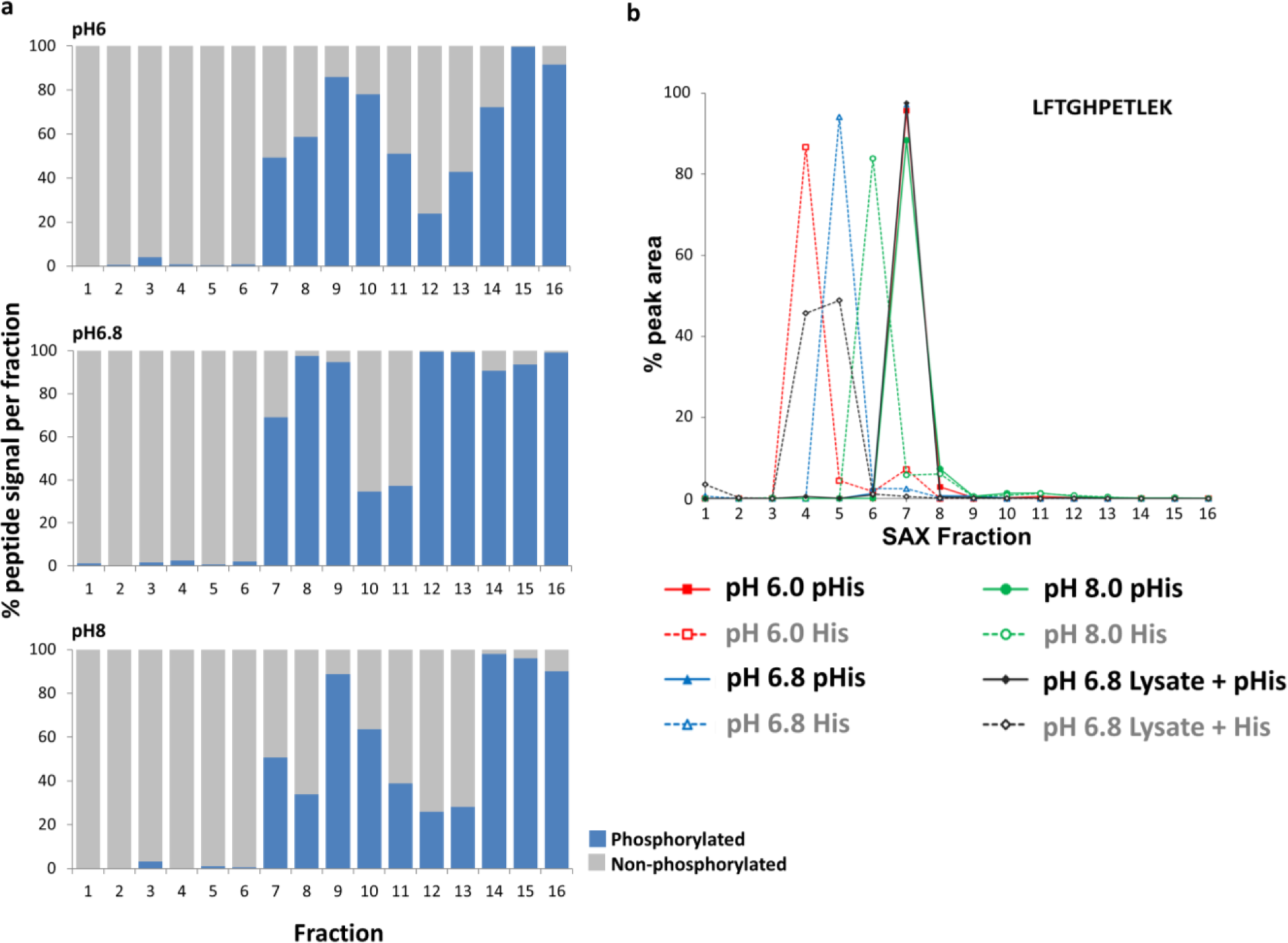
Optimal separation of pH is myoglobin peptides from non-phosphorylated counterparts by strong anion exchange (SAX) is achieved at pH 6.8. **(a)** Percentage of the total signal intensity attributed to phosphopeptides (blue bars) and non-phosphorylated peptides (grey bars) following SAX fractionation at pH 6.0, pH 6.8 and pH 8.0, with later fractions consisting of up to 100% phosphopeptide ion signal. **(b)** Representative pHis-containing myoglobin peptide (LFTG<u>H</u>PETLEK; solid line), and its non-phosphorylated counterpart (LFTGHPETLEK; dashed line), quantified across all 16 SAX fractions; SAX was performed at pH 6.0 (red), pH 6.8 (blue) or pH 8.0 (green). % of the total peak area of each individual peptide across the gradient is plotted for each fraction. In order to assess pHis stability and effects of SAX separation in a complex mixture (black), SAX was also repeated at pH 6.8 with phosphorylated myoglobin spiked into a human cell lysate prior to digestion. Data for the five pHis-containing myoglobin peptides is shown in Supp. Fig. 5.

### Precursor neutral loss ions do not improve confidence in identification of pHis-containing peptides

It has previously been reported that pHis-peptides exhibit a unique gas-phase fragmentation ‘fingerprint’ when subjected to collision-induced dissociation (CID), arising as a result of neutral loss of 80, 98 and 116 atomic mass units (amu) from the precursor ion^40^. Consequently, this triplet neutral loss pattern could theoretically be exploited to improve identification of pHis-containing peptides from LC-MS/MS data. In agreement with these findings, we observed characteristic triplet neutral losses in the tandem mass spectra of all five pHis myoglobin peptides following both ion-trap CID and higher energy collision dissociation (HCD), which represents the optimal fragmentation regime for phosphopeptide identification using the Orbitrap platform^41^. To investigate the utility of triplet neutral loss as a potential signature for accurate pHis site localisation and high-throughput pHis characterisation in complex samples, we assessed precursor neutral loss for phosphopeptides with confident phosphosite localisation (1% FLR) derived from a human HeLa cell lysate known to contain pSer, pThr, pTyr and pHis, using either CID or HCD-based fragmentation (Fig. 2). Of the 236 (CID) or 342 (HCD) singly phosphorylated peptides identified from a single SAX fraction, Fig. 2a), 5% or 3% (CID or HCD respectively) contained all three neutral loss ions when considering all phosphopeptides with *ptm*RS score >75%. As previously reported,^41^ significant numbers (24% by CID, 49% by HCD) of phosphopeptides generated no precursor ion neutral loss fragments, considering all ions with signal intensity ≥5% of the base peak, with little improvement when considering all ions ≥2% relative signal intensity (17% for CID, 30% for HCD). We subsequently evaluated the prevalence of precursor neutral loss by HCD or CID for each category of phosphopeptide according to the type of residue modified (Fig. 2b,c). CID spectra of pTyr peptides exclusively exhibited no, or a single neutral loss ion, while 30% of the equivalent HCD spectra contained two or more neutral loss ions. Crucially, none of the endogenous singly phosphorylated pHis-containing peptides identified in this SAX fraction exhibited triplet neutral loss by either CID or HCD (Fig. 2b,c).

The prevalence of HCD-induced precursor neutral loss for unique singly phosphorylated peptides containing either pHis, pSer/pThr, or pTyr was considered across all SAX fractions (Fig. 2d), as a function of site localisation confidence (*ptm*RS score >99%, 75-99%, 25-75%, <25%). Although a small proportion of confidently localised pHis peptide HCD spectra (*ptm*RS >75%) were observed with the triplet pattern across all the SAX fractions (^~^8%), phosphopeptides modified on other residues also exhibited a similar or greater proportion of triplet neutral loss, with 7% of spectra from pSer/Thr-containing peptides (*ptm*RS>99%) containing all three neutral loss ions. Manual examination of spectra confirmed that absence of this triplet pattern, or indeed any neutral loss following HCD, was not sufficient to rule out pHis-peptide identification. It was noticeable that indeed, the majority (>70%) of phosphopeptides containing confidently localised pHis or pTyr (*ptm*RS>99%), were observed with no neutral loss, although this was not as marked for pSer/Thr-containing peptides, where only 30% of peptides across all SAX fractions exhibited no neutral loss. These findings agree with previous observations demonstrating that phosphopeptide neutral loss propensity is dependent on both the type and relative position of amino acids within the sequence^42-44^.

Further examination of the neutral loss ions (Δ80 amu *vs* Δ98 amu *vs* Δ116 amu) revealed that the nature of the phosphorylated residue had very little effect on the likelihood of observation of a Δ80 amu ion from the precursor following HCD (Fig. 2e-g). As expected, neutral loss from pSer/pThr peptides was predominantly 98 amu, often with additional loss of H_2_O (Δ116 amu). Consequently, pSer/Thr-containing peptides were much more likely to generate spectra with Δ116 amu than either pTyr or pHis-containing peptides. Interestingly, the neutral loss pattern from pTyr-containing peptides was broadly similar to that observed for pHis peptides, suggesting similar mechanisms for neutral loss. Although the data presented only considers ions at >5% base peak intensity, there was no significant difference at either 2% or 10% relative intensity (Supp. Figs. 6 and 7). These data suggest that neither the presence, nor absence, of the triplet neutral loss ‘fingerprint’ can conclusively define the type of phosphorylated residue present.

### Comparative analysis of the human (phospho)proteome following PHPT1 knockdown

To evaluate UPAX for global pHis-peptide identification, and with the intention of increasing ‘basal’ levels of histidine phosphorylation, we employed siRNA in HeLa cells to reduce expression of one of the only two suspected mammalian phosphohistidine phosphatases, PHPT1/PHP14^45, 46^ prior to SAX fractionation and phosphoproteome analysis (Fig. 3a). Western blotting confirmed efficient knockdown of PHPT1 (Fig. 3b), and sixteen SAX fractions were subsequently collected and analysed by LC-MS/MS using HCD nlEThcD. A representative UV trace for the SAX separation (Fig. 3c), and associated base peak chromatograms for three of the 16 SAX fractions are shown (Fig. 3d–f). All MS/MS data analysis was performed using Proteome Discover, invoking MASCOT for peptide identification and *ptm*RS to evaluate phosphosite localisation confidence (Fig. 3g). To assess differential protein regulation between control (non-targeting siRNA; NT) and PHPT1 knockdown samples, proteins identified in two or more bioreplicates were taken forward. 83% of the 4580 unique proteins identified across the two conditions were common, indicating a high degree of overlap (Supp. Fig. 8). Of those protein identifications that were condition specific, there was no discernible difference in gene ontology analysis with respect to biological processes, molecular function or protein class (Supp. Fig. 9), suggesting limited functional consequences of PHPT1 depletion. Considering non-phosphorylated peptides observed at least twice across the dataset, 83% of those from the PHPT1 depleted sample were also identified in the control lysate. This overlap reduced to 76% when considering phosphopeptides containing pHis, pSer, pThr or pTyr. Independent of assessment criteria, slightly greater numbers of phosphopeptides were consistently identified in control (NT) siRNA *versus* PHPT1 siRNA samples.

**Figuire 2.**
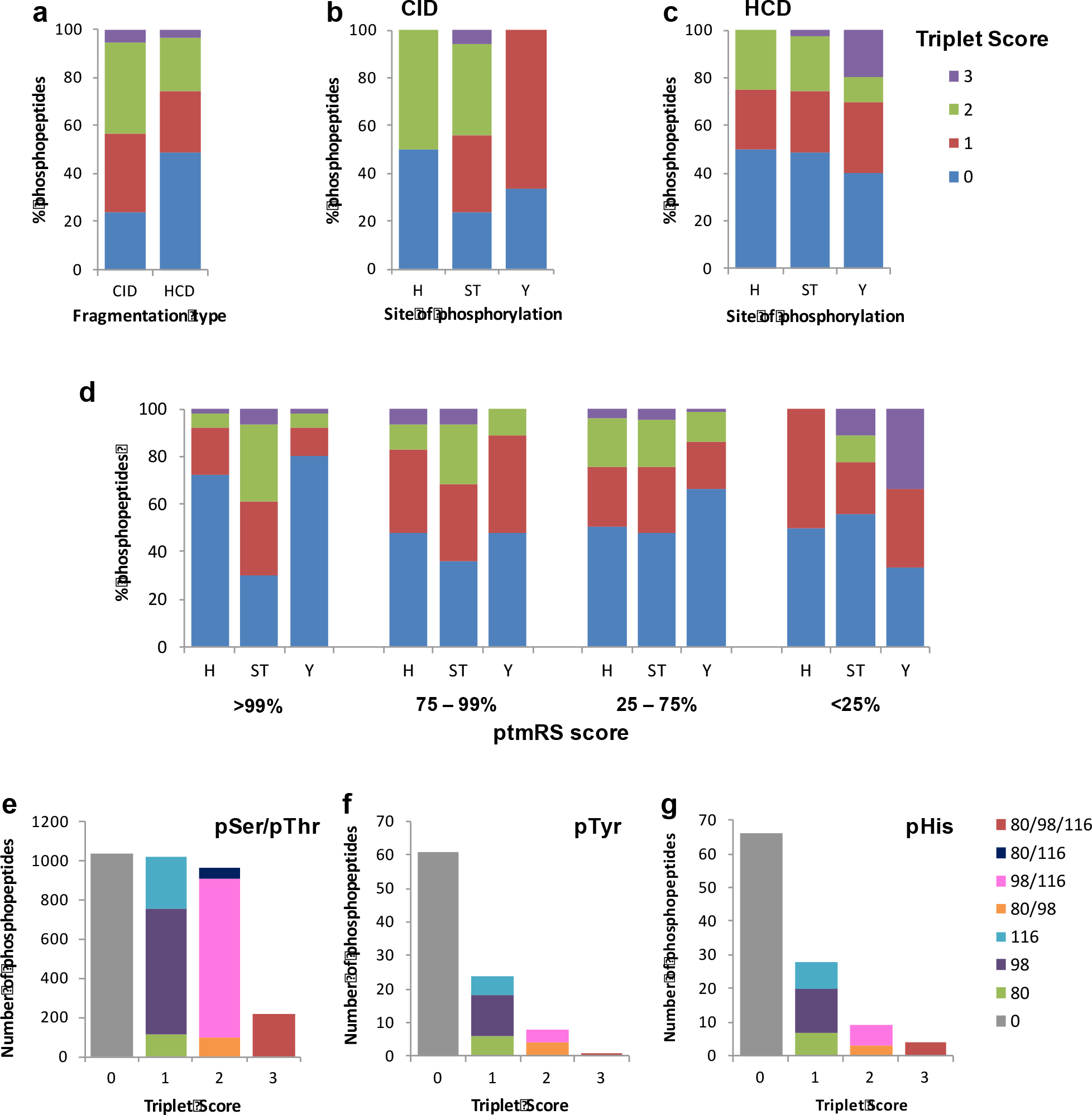
Phosphopeptide neutral loss pattern is not diagnostic for histidine phosphorylation. (**a**) Comparison of triplet score following either CID or HCD from unique HeLa cell derived phosphopeptides with ptmRS score>75%. (**b, c**) Evaluation of triplet scores for unique singly phosphorylated pHis (H), pSer/pThr (ST) or pTyr (Y)-containing peptides (ptmRS score>75%) following either (**b**) CID or (**c**) HCD. All data are from SAX fraction 13. (**d**) Distribution of triplet score as a function of site localisation confidence (ptmRS score) for unique singly phosphorylated peptides across all SAX fractions. (**e-g**) Numbers of phosphopeptides (ptmRS score>75%) exhibiting different combinations of the three neutral loss species (Δ80, Δ98 and/or Δ116 amu from precursor at >5% relative signal intensity to base peak) for pSer/pThr (e) or pTyr (f) or pHis (g). Triplet score of: 3 indicates observation of all three neutral loss ions; 2 indicates any two neutral loss species; 1 indicates any one of the three neutral loss ions.

**Figure 3.**
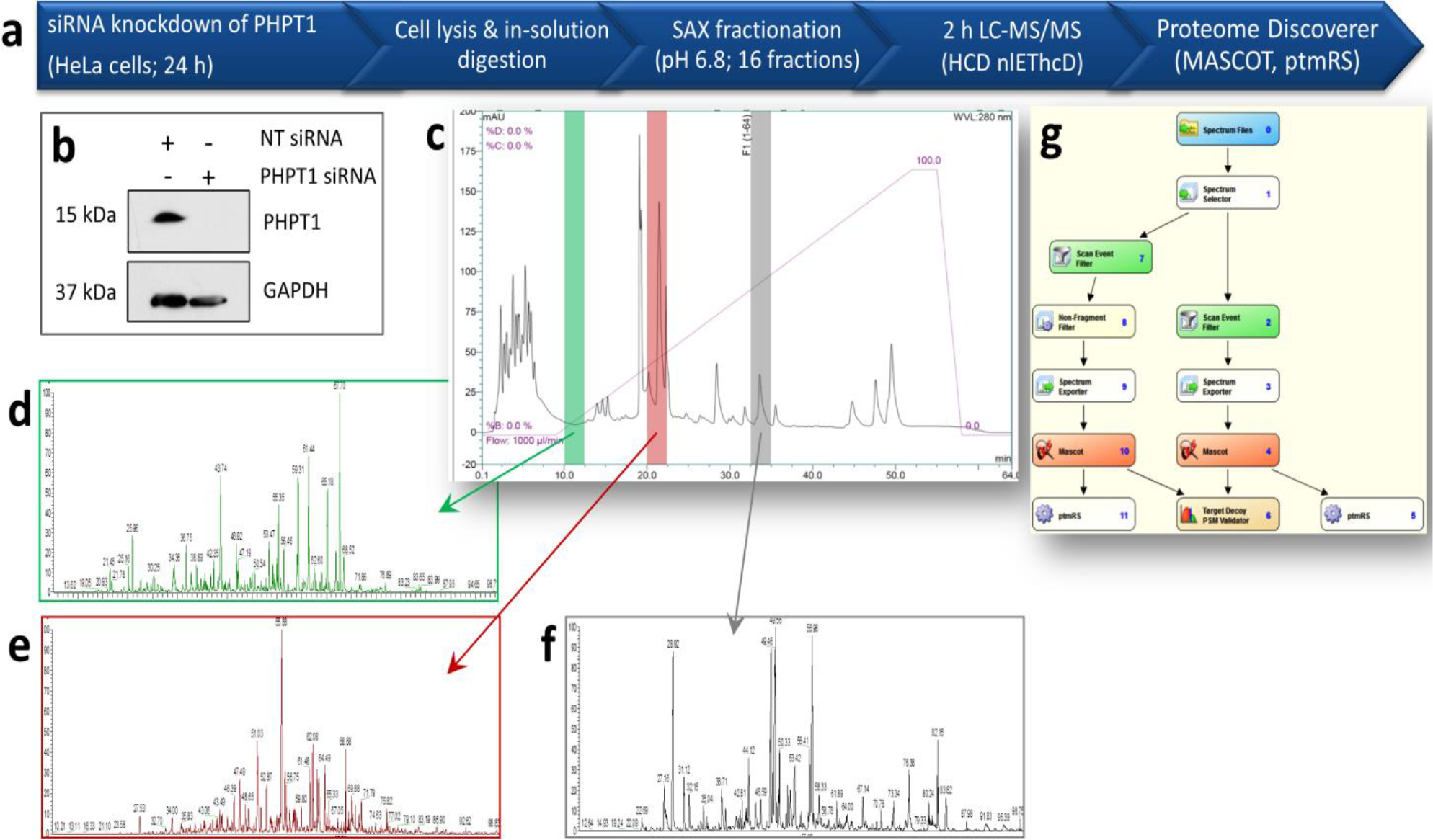
Workflow for unbiased phosphoproteomics. (**a**) Schematic representation of the UPAX strategy for phosphopeptide enrichment and analysis, including of phosphohistidine-containing peptides. (**b**) siRNA-mediated knockdown of PHPT1 in HeLa cells (48 h) with reference to GAPDH loading control. NT - non-targeting siRNA control. (**c**) Representative SAX profile of trypsin digested HeLa lysate (Abs_280 nm_). Base peak chromatograms are shown for select SAX fractions following high-resolution LC-MS/MS using an Orbitrap Fusion mass spectrometer: (**d**) fraction 3 (green); **(e)** fraction 6 (red); (**f**) fraction 10 (grey). Peptides were fragmented by HCD, with neutral loss of 98 amu from the precursor ions triggering EThCD for improved site localisation confidence (Ferries et al.). **(g)** Proteome Discoverer pipeline for data analysis. Tandem mass spectra were separated according to fragmentation strategy prior to searching with Mascot. The ptmRS node was used for phosphosite localisation. Analysis was performed on three independent biological replicates.

### Distribution of cellular pHis and phosphorylation motif characterisation

A total of 310 unique novel sites of histidine phosphorylation were identified across all samples (NT and PHPT1 siRNA) at a 1% FLR (Fig. 4a). These were distributed across 299 peptides derived from 294 proteins (Supp. Table 5). Of these proteins, <10% had previously been reported to be phosphorylated on His, by virtue of specific immunoprecipitation with monoclonal antibodies against 1- or 3-pHis-containing proteins^12^ (Supp. Table 6), and we did not identify any of the limited number of pHis sites previously reported in human cell extracts^47^.

**Figure 4.**
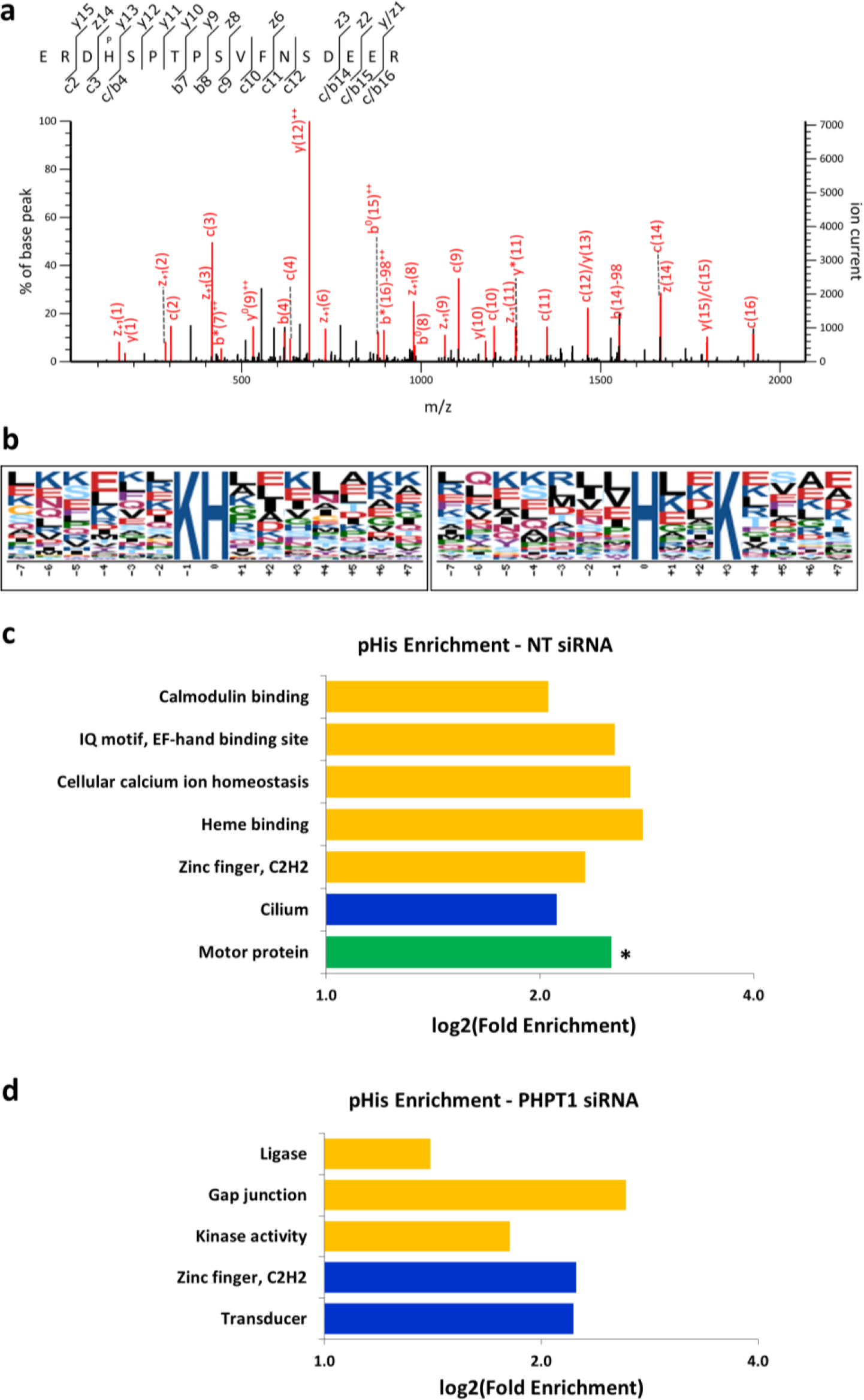
Amino acid motif analysis and categorisation of phosphohistidine-containing proteins. (**a**) Annotated EThcD product ion mass spectrum of the triply protonated peptide ion at m/z 694.623 representative of ERDpHSPTPSVFNSDEER from FIP1_HUMAN. (**b**) Motif analysis of the residues (+/-5) surrounding the ptmRS defined sites of pHis from all identified pHis-containing peptides. (**c, d**) Functional annotation clustering of pHis-containing proteins using DAVID, following treatment of HeLa cells with either non-targeting (NT) siRNA (**c**) or PHPT1 siRNA (**d**) prior to phosphoproteomics analysis. Only those classifications represented by >3 proteins are depicted. Colours are indicative of P-value: P<0.001 (green); P<0.01 (blue); P<0.05 (yellow). Asterisks define those clusters where the Benjamini-Hochberg adjusted P-value<0.1. See supplementary tables for details.

Our analysis represents the first large-scale *unbiased* study to pinpoint specific sites of pHis in any organism. Consequently, we examined amino acid conservation surrounding the sites of His phosphorylation. Combining all pHis-containing peptides where pHis was confidently localised (1% FLR), we observed two pHis-based motifs using Motif-X^48, 49^. Both motifs were Lys-directed, with Lys enriched at either -1 (2.85 fold) or +3 (2.58 fold) positions (Fig. 4b). There was also a slight preference for Lys, Glu and hydrophobic residues, in particular Leu, in the ^~^5 residues surrounding pHis. Unsurprisingly given the charge-directed nature of SAX, there was a bias in the motifs observed in the early versus later eluting ion exchange fractions (Fig. 4b; Supp. Fig. 9, Supp. Table 7), with the later eluting fractions (9-16) exhibiting strong preference for Glu at +5.

PANTHER-based gene ontology enrichment analysis of all confidently localised pHis-containing proteins compared with all proteins identified in the samples, revealed a 17-fold increase in ‘dynein light chain binding’ (P=2.79E-2), including dynein heavy chains (5 and 14) and cytoplasmic dynein 1 and 2 heavy chain 1. In parallel, the dynein complex was upregulated as a cellular component (12-fold; P=1.66E-2). A number of proteins classified also exhibit ATP-dependent microtubule motor activity, thus this molecular function was also upregulated (P=2.33E-3). General ATPases were upregulated ^~^4 fold (P=2.99E-2), including the DNA replication licensing factor MCM6, the RNA helicase DDX42 and several cation transporters. These classifications were not significantly upregulated in the PHPT1 siRNA samples, suggesting a lack of general pHis modulation by this phosphatase.

Consistent with these findings, independent analysis with DAVID (v6.8)^50, 51^ (Fig. 4 c, d; Supp. Fig. 10 a, b, Supp. Table 8) identified outer dynein arm assembly (P=3.3E-3) and microtubule motor activity (P=1.3E-3) as enriched for pHis, as were ciliopathy and cilium (P= 7.2E-3), haem binders (P=2.7E-2) and cation transporting P-type ATPase (P=3.8E-2). No KEGG pathways were significantly enriched in the control datasets. In contrast, G-protein coupled receptors were enriched (P=4.2E-3) in the PHPT1 siRNA samples, as was cAMP signalling and the GAP junction pathway, by virtue of five (P=1.9E-2) and four associated proteins respectively (P=2.5E-2).

### Phosphorylation of Lys, Arg, Asp and Glu amino acids is readily detected in human cell extracts

Like pHis, phosphorylated Asp, Lys, Arg, Glu are acid-labile^14, 15^. Since our UPAX workflow is performed at near physiological pH, we searched our MS datasets for other acid-labile phosphopeptides. Given our observation that numerous pHis-containing peptides were co-phosphorylated on Ser, Thr or Tyr (Supp. Table 5), searches for non-canonical sites of phosphorylation were performed with the option of additional Ser, Thr or Tyr phosphorylation. Remarkably, in addition to ^~^300 pHis-containing peptides, a significant number of peptides were found to contain distinct non-canonical phosphorylated residues.

In total, we identified 2740 phosphopeptides where a site of modification could be confidently localised to amino acids other than Ser, Thr, Tyr or His (Fig. 5a), including phosphorylated Arg, Lys, Asp and Glu. No Cys phosphorylated peptides were identified in these samples. Evaluation of the relative numbers of phosphosites identified for each of these residues confirms that observation of a pX site is not purely a consequence of an increased prevalence of this individual residue (Fig. 5b). Indeed, although the total number of Glu residues in our dataset was almost equal to that of Ser, only one-tenth the number of pGlu sites were identified compared to pSer. To understand the distribution and potential roles of individual non-canonical phosphosites, we employed informatics analysis of phosphorylated proteins using DAVID and PANTHER.

**Figure 5.**
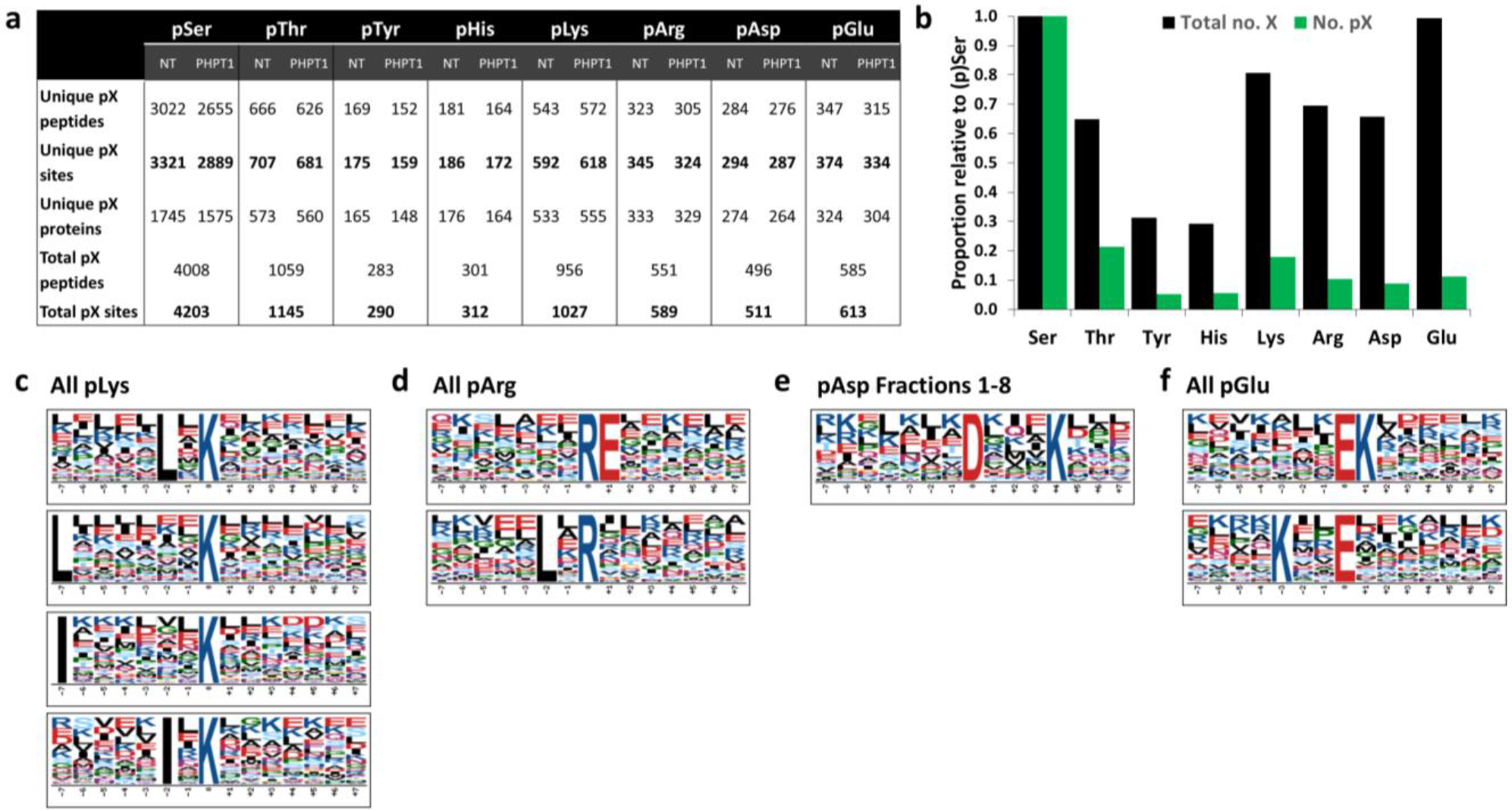
Identification of acid-labile phosphorylation sites. (**a**) Numbers of unique phosphopeptides and phosphosites identified with phosphorylation of Ser, Thr, Tyr, His, Lys, Arg, Asp or Glu in HeLa cell extracts treated with non-targeting (NT) siRNA, or PHPT1 siRNA. (**b**) Relative proportion of the total number of modifiable residues (black) in the identified proteins, and the number of phosphorylated sites identified for that residue (green) relative to Ser or pSer respectively. Motif analysis of the residues (+/-5) surrounding the ptmRS defined sites of phosphorylation for (**c**) pLys, (**d**) pArg, (**e**) pAsp, (**f**) pGlu. With the exception of pAsp, all identified pX containing peptides were used. Motif defined for pAsp, only used phosphopeptides present in SAX fraction 1-8.

### Phospholysine

Even though phospholysine (pLys) was demonstrated to be present at significant levels in rat liver ^~^50 years ago^52-55^, and a phosphatase with *in vitro* pLys phosphatase activity has been reported^56^, very little is known about the prevalence or positional distribution of this modification. pLys is not currently considered in the controlled vocabulary used in UniProtKB and consequently no pLys sites of modification have been recorded in any organism. Of all the non-canonical phosphorylation sites identified in our analysis, pLys was the most abundant in relative terms, with 592 pLys sites identified under ‘basal’ conditions and 618 pLys sites recovered after PHPT1 knockdown (Fig. 5a; Supp. Table 9). Motif analysis indicated a marked preference for Leu or Ile at -2 or -7 positions and some enrichment for non-phosphorylated charged amino acids (Asp, Glu, Lys and Arg) upstream and downstream of the site of Lys phosphorylation (Fig. 5d; Supp. Fig. 11).

The increase in the number of pLys peptides identified in the PHPT1 siRNA samples, which is in contrast to the slight decrease seen with other phosphorylated amino acids analysed (Figure(Fig. 5a), hints that PHPT1 may regulate pLys turnover in cells, and is consistent with the ability of PHPT1 to directly dephosphorylate pLys-containing proteins *in vitro^56^*.

PANTHER analysis revealed that pLys-containing proteins associated with the cytoskeleton were enriched 1.5 fold (P=4.82E-03) with respect to all proteins identified, as were (intracellular) non-membrane bound organelles (1.3 fold, P=1.76E-2), while cytoplasmic part and mitochondrion were both down-regulated for pLys (0.8 fold, P=1.79E-04; 0.5 fold, P=5.32E-03 respectively). Significant enrichment was also observed for ‘plasma membrane-bound cell projection’ proteins after PHPT1 knockdown (1.8 fold, P=2.04E-2). DAVID analysis also indicated significant enrichment for pLys in cell junction (total of 35 proteins; P=5.1E-4) and motor proteins (P=5.2E-5) (Supp. Fig. 10 c, d, Supp. Table 10). Calponin homology domain (P=9E-4), cadherin binding involved in cell-cell junction (P=9.1E-3), ankyrin repeat (P=1.3E-2), and PDZ-domain containing proteins (P=2.8E-3) were also enriched, with 15 proteins defined in the latter category including tight junction protein 1/2, AHNAK nucleoprotein, and PP1 regulatory subunit 9A. There was also a small (^~^1.4-fold), but significant, enrichment in ATP-binding (P=5.2E-3 for control; P=9.6E-5 for PHPT1 siRNA) and a 2.4-fold enrichment for ATPase activity (P=7.8E-4). Following depletion of PHPT1, IQ motif-containing proteins with EF-hand binding sites (15 proteins; P=1.1E-6) was specifically enriched, including a number of muscle proteins (P= 5.36E-04) also found to be enriched under basal conditions. Five transcription factors containing the AT-rich interaction (ARID) DNA-binding domain (P=3.8E-3) were also upregulated 7-fold after depletion of PHPT1.

Importantly, twelve KEGG pathways were significantly enriched for pLys-containing proteins (P<5E-2; Supp. Table 10, Supp. Fig. 10 c, d), most notably tight junction (12 proteins, including myosins; P=1.7E-3), feeding primarily into cell survival/differentiation, and the oxytocin signalling pathway (11 proteins; P=6.5E-3). The tight junction is also one of the 11 KEGG pathways enriched following PHPT1 knockdown (13 proteins; P=1.0E-3), although the oxytocin signalling pathway was not. Instead, PHPT1 siRNA induced a 3-fold enrichment in the ‘prostate cancer’ pathway, as represented by 9 known constituents.

### Phosphoarginine

Arginine kinase activity has been described in vertebrates. The highly basic proteins histone H3 and H4, and myelin basic protein (MBP), both of which are promiscuous protein kinase substrates, are the only substrates reported^57-59^. In our analysis, the number of human pArg sites identified was almost double that identified for pHis, with 345 and 324 unique pArg sites catalogued before and after PHPT1 knockdown (Fig. 5a, Supp. Table 11). Interestingly, two distinct consensus motifs were identified for pArg, with either a charged Glu placed at +1, or a hydrophobic Leu positioned at -2 with respect to the site of Arg phosphorylation (Fig. 5e; Supp. Fig. 12).

Specifically, voltage-gated channels (4.5 fold change, P=2.3E-2), collagen (4.4 fold change, P=2.6E-2), and tetratricopeptide (TPR) repeat-containing proteins (2.1 fold enrichment, P=2.12E-02) were all significantly enriched for pArg under basal conditions (Supp. Fig. 10 e, f; Supp. Table 12). Following PHPT1 depletion, there was significant enrichment in ATP binding proteins (48 proteins; P=7E-4), including 11.6 fold upregulation of dynein-related proteins exhibiting ATPase activity (P=4.3E-3) and multiple members of the ATP binding cassette subfamily. ABC transporter-like (P=1.1E-2), spectrin/alpha-actinin repeat (P=3.9E-3), including AKAP6, dystrophin, periplasm, and calmodulin binding (P=3.4E-2) are also enriched ^~^3 fold.

A four-fold enrichment was also observed in the KEGG mTOR signalling pathway (P=3.1E-2), ^~^2 fold enrichment in PI3-Akt signalling pathway (11 proteins identified, P=3.5E-2) and ^~^3-fold enrichment in the TNFα signalling pathway (P=4.6E-2). Interestingly, following PHPT1 depletion, cancer-associated focal adhesion (P=3.3E-2) and RAS signalling pathways were also selectively enriched (Supp. Fig. 10 e, f; Supp. Table 12).

### Phosphoaspartate

Although phosphorylated Asp is recognised as an intermediate in enzyme catalysed reactions (*e.g*. P type ATPases, HAD phosphatases, atypical RIO2 and myosin heavy chain A kinases and phosphomutases^14, 60-62^) we believe that the 292 unique sites (274 distinct proteins) from control samples, and the 284 pAsp sites (264 proteins) observed here after PHPT1 depletion, are novel (Supp. Table 13). Interestingly, RIO1 kinase was observed to contain pAsp in the control dataset, although the site of modification, pAsp20, is outside of the catalytic domain (Supp. Table 13) in contrast to that observed for RIO2. Searching UniProtKB for 4-aspartylphosphate as a modification type in *homo sapiens* did not identify any proteins known to contain pAsp. No common consensus motif for pAsp was observed when considering our phosphosite dataset as a whole. However, Motif-X analysis of early eluting SAX fractions (1-8) suggested a preference for pAsp when Lys was installed at position +4 (Fig. 5c; Supp. Fig. 13). This is distinct to a pDXXX(T/V) motif previously reported in ATPases^63, 64^, although a pAsp in a DXXXV motif was identified in one of the ATPases in our dataset (Supp. Table 13).

To reveal putative functions of pAsp in human cells, we performed bioinformatics analysis (Supp. Table 14, Supp. Fig. 10 g, h). TPR-containing domains were enriched in control and PHPT1 siRNA samples (P=1.2E-6, P=4.5E-3 respectively) with ^~^14 proteins represented. Given the involvement of this domain in regulating assembly of multiprotein complexes, a role for pAsp in regulating complex formation is hypothesised. In control samples, cilium-resident proteins were also enriched (P=1.9E-3), as were microtubule (P=7.1E-3), cell junction (P=2.6E-3) and cell cycle components (P=2.8E-3) (Supp. Table 14, Supp. Fig. 10 g, h). Interestingly, PHPT1 depletion led to increased pAsp-containing proteins amongst ATP-binding (P=3.5E-4) and ATPase activity (P=5.5E-4) proteins, with 46 and 13 respective proteins in these classifications. Helicases (P=6.73E-03) and motor proteins (P=6.60E-03) were also upregulated ^~^3 fold in response to PHPT1 depletion (Supp. Table 14, Supp. Fig. 10 g, h). No KEGG pathways were significantly enriched in either sample set.

### Phosphoglutamate

Under control conditions, 375 unique pGlu sites were identified across 347 peptides, decreasing to 284 following PHPT1 knockdown (267 phosphopeptides), of which 77 were identified under both experimental conditions (Supp. Table 15). Like pLys and pAsp, pGlu is not listed in UniProt as a defined modification, thus there is no curated record of endogenous modification of proteins on this residue. In the free-form, pGlu is known as gamma glutamyl phosphate, which is an intermediate during the glutamine synthase-mediated conversion of glutamate to glutamine. However, very few proteins are established to be modified by phosphorylation on Glu^14^. Motif analysis of all pGlu-containing peptides indicated a strong preference for Lys at +1 or -3 (Fig. 5f; Supp. Fig. 14), suggesting previously unsuspected signalling functions. Consistently, >18-fold enrichment was observed for pGlu proteins amongst dynein heavy chain proteins (P= 8.79E-05). GTPase activation (3-fold enrichment; P=1.34E-03), fibronectin proteins (4.4 fold enrichment; P=2.11E-03), and ATP binding proteins (P= 8.03E-03) were also enriched for pGlu (Supp Fig. 10 i, j; Supp. Table. 16). Although ATP binding proteins were observed under both conditions, ATPase activity was notably enriched after PHPT1 knockdown, suggesting biological significance of PHPT1 in regulating Glu phosphorylation. Platelet activation was the most significantly enriched KEGG pathway (^~^3-fold; P=4.30E-02), but only following PHPT1 depletion (Supp. Fig. 10 i, j; Supp. Table. 16).

### A putative biological role for His phosphorylation in protein kinase regulation

The reversible regulation of eukaryotic protein kinases (ePKs) by PTMs, most notably Ser, Thr and Tyr phosphorylation, is a central eukaryotic signalling mechanism. Of the ^~^300 pHis-containing peptides that we identified, 11 (^~^4%) were found in human protein kinases, ^~^2-fold higher than expected for the ePK superfamily by chance alone. Furthermore, three pHis-derived peptides (identified multiple times across samples) mapping to the Ser/Thr kinases PLK2 and CDK19 and the tyrosine kinase Ephrin A1 (Supp. Table 5), were localised to an identical region in the catalytic domain, specifically the HxN-His motif of the αC-β4 loop, which is implicated in positioning of the αC helix and Hsp90-mediated kinase maturation^65, 66^ (Fig. 6). Interestingly, a His residue is conserved in >65% of human protein kinases, and is a phosphorylatable Ser/Thr in a further 10% of ePKs found across the various taxa. Another His phosphorylation site is found in the tyrosine kinase KIT (pHis630) adjacent to the β3 loop. PTM of the αC-β4 loop has been observed in multiple kinases and is proposed to regulate the position of the preceding αC-helix^67, 68^. Consequently, we hypothesised that phosphorylation of the HxN-His residue might regulate protein kinase activity by altering protein dynamics.

We therefore performed molecular dynamics (MD) simulations on four possible pHis139 states of PLK2 based on the protonation state and attachment of the phosphate group in this histidine (pH139-N1-H1D, pH139-N1-H2D, pH139-N3-H1E, pH139-N3-H2E) (See Methods). Comparison of these simulations suggests increased flexibility in the phosphorylated forms compared to the non-phosphorylated form of PLK2 (Fig. 6b). In particular, the 3-pHis (N3) form displays increased flexibility in the activation segment; the regulatory C-helix, the catalytic HRD motif (Fig. 6b, 6c) and the regulatory spine residue (H134) also undergo a conformational change (Fig. 6d).

**Figure 6.**
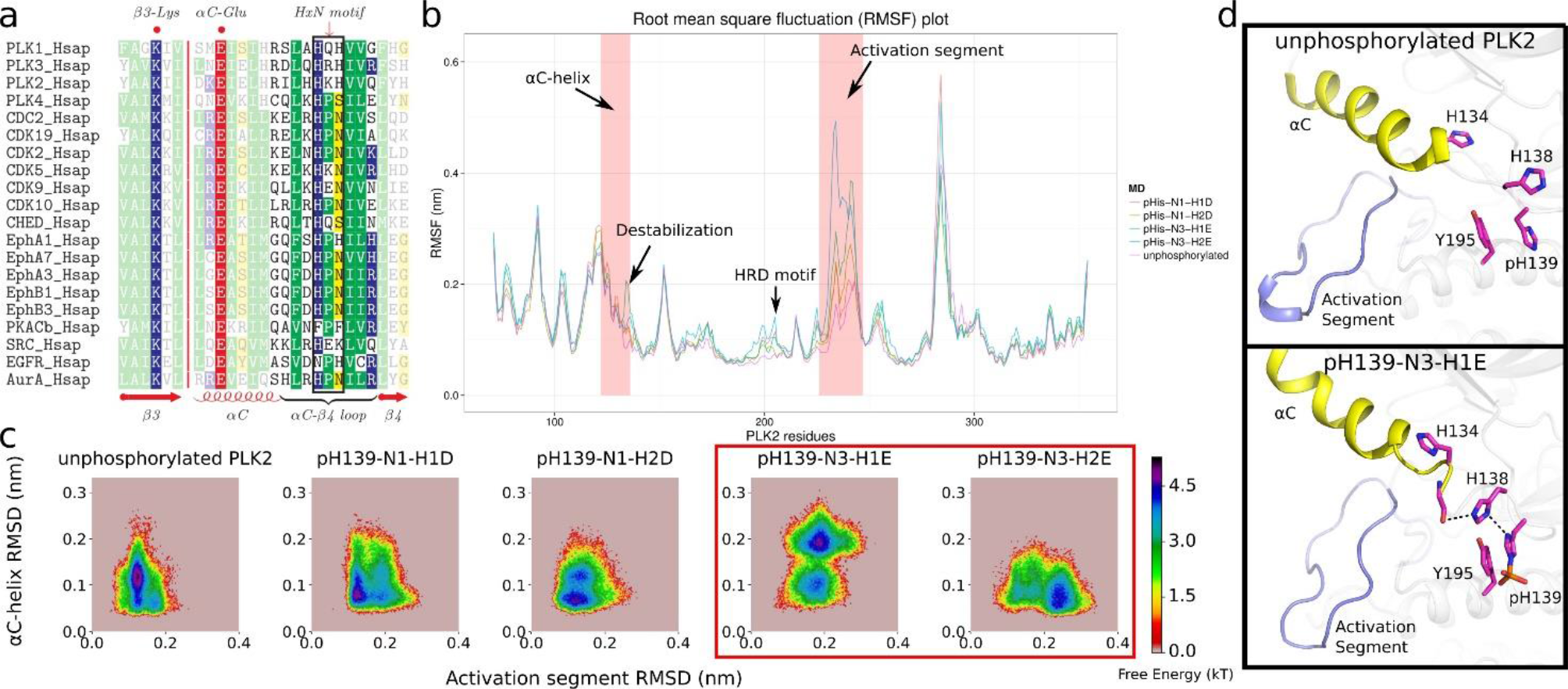
Molecular dynamics simulations of PLK2 with H139 phosphorylation. (**a**) Alignment of selected human kinases highlighting the HxN motif. (**b**) Root mean square fluctuation (RMSF) plot of non-phosphorylated PLK2, pH139-N1-H1D, pH139-N1-H2D, pH139-N3-H1E, pH139-N3-H2E (See methods). Key regions that display differential fluctuations are shown. (**c**) Free energy landscape through the root mean square deviation (RMSD) of αC-helix and activation segment. (**d**) Representative snapshot of non-phosphorylated PLK2 (upper panel) and pH139-N3-H1E (lower panel).

## Discussion

In this paper, we describe an experimental pipeline suitable for identification of both canonical (pSer, pThr, pTyr) *and* non-canonical (pHis, pAsp, pGlu, pLys, pArg) phosphorylation sites from complex biological samples, which we have validated with human cell extracts. Using this approach, termed UPAX, we reveal extensive phosphorylation of acid-labile phosphopeptides derived from human proteins. We pinpointed over 1700 novel non-canonical phosphosites under standard cell culture conditions, and defined several conserved motifs surrounding these sites of phosphorylation. The five phosphorylated acid-labile conjugates identified (His, Asp, Glu, Arg and Lys) occur on amino acids that together account for ^~^25% of the total number of residues found in typical vertebrate proteins, and this number increases to over 40% if non-canonical Ser, Thr and Tyr residues are included. The prevalence of non-canonical phosphorylation, and in particular our finding that pLys exceeds that of pThr in the samples analysed, was surprising. In addition, we predicted that knockdown of PHPT1, one of only two previously described mammalian His phosphatases, might increase the global levels of pHis. Instead, we measured a marked increase in pLys, but not pHis, peptides present under these conditions, suggesting that PHPT1 may actually function as a rate-limiting cellular pLys phosphatase, consistent with an ability to dephosphorylate pLys *in vitro*. Pathway analysis did not reveal clustering of acid-labile PTMs in any particular sub-cellular compartment, with the exception of pLys, which was enriched in the cytoskeleton and downregulated in both cytoplasm and mitochondrion. These findings point towards generalised functions of these phosphorylated residues in human cell biology.

Despite the extensive non-canonical phosphorylation revealed in this study, high throughput phosphopeptide analysis by MS presents experimental challenges. In particular, the negative free energy of phosphoramidate bond hydrolysis in pHis/pLys/pArg means that peptides containing these non-canonical phosphosites are considered more likely to undergo phosphate scrambling than the relatively low mobility reported amongst canonical phosphopeptides^69^. The potential for phosphate transfer from non-canonical phosphopeptides during collision-mediated fragmentation is already established for various model peptides ^25, 70-72^. Importantly, however, careful analysis of pHis myoglobin standards spiked into cell lysates did not reveal positional phosphate transfer of any of the five phosphopeptides with either HCD or ET-mediated dissociation (ETcaD or EThcD). However, we cannot currently rule out that a proportion of the phosphosites reported here may be due to HCD-mediated phosphate re-arrangement. For example, pArg has been reported to translocate to Ser and Glu following HCD, resulting in <20% of the pArg sites in a predefined sample being incorrectly localised^72^, and we have previously observed intermolecular transfer of pHis during low energy CID, albeit at high peptide concentrations^25^. Application of ETD, rather than HCD, fragmentation could potentially overcome this type of phosphate translocation. However, ETD is inefficient for doubly-charged peptide ions and, dependent on precursor ion charge state and amino acid composition, phosphate scrambling has also been observed for pLys-containing peptides during ETD^73^. In terms of our findings, the transfer of phosphate from N-linked donors could theoretically reduce the total numbers of pHis, pLys, pArg sites identified, with a concomitant increase in peptides containing, most likely, pSer.

To reduce the unnecessarily large search space associated with simultaneously considering 8 potential phosphosites, we searched for each of the five distinct pX residues in combination with variable pSer, pThr and/or pTyr. This strategy also minimised the potential for enhanced false positive phosphopeptide identification. However, given that only a single non-canonical pX site is considered at any one time, there is potential for incorrect site assignment, as is the case whenever specific variable modifications are defined, rather than performing an ‘open modification’ search. Indeed, some 23% of the 3237 total peptide spectrum matches (PSMs) containing non-canonical phosphorylation across all searches were derived from the same MS/MS scan, with a different pX site defined at a 1% FLR for the same peptide sequence. Two possible scenarios together explain these results: i) the site of phosphorylation was incorrectly assigned in one of the searches simply because the correct pX site was not offered as a potential site of modification, or ii) both forms of the phosphopeptide are present in the sample. In the majority of cases where the pX site was differentially localised, the putative sites of phosphorylation are in close proximity (<^~^3 residues), suggesting that phosphosite assignment is ambiguous due to insufficient site localisation-determining product ions. Open modification searches, such as MS Amanda^74^ and MSFragger^75^, could potentially be used to discriminate these possibilities, but unlike *ptmRS* which we have used here, neither are currently benchmarked for phosphosite localisation confidence.

The general non-random nature of His, Arg, Lys, Asp and Glu phosphorylation site deposition on cellular proteins suggests a myriad of undiscovered biological roles for these modifications, which potentially rival those being catalogued for pSer, pThr and pTyr. In terms of mechanism, the introduction of a negatively charged phosphate group adjacent to the basic side chain of His, Arg and Lys residues are of particular interest. The addition of one (or more) phosphate groups not only changes the size and polarization of the side chain, but also reverses local net charge. By virtue of the near neutral pKa value for the imidazole proton, His rapidly undergoes catalytically important transitions from a charged to uncharged side-chain in many proteins. Consequently, phosphorylation might have a major impact on enzyme-mediated catalysis as well as acting as a conduit to couple (reversible) phosphorylation with modular cellular signalling in a manner reminiscent of well characterised pTyr:SH2/PTB domain interactions of vertebrates^76^. The acidic side chains of Arg and Lys are assumed to be fully protonated under physiological conditions, and the introduction of a phosphate group with a new directional net negative charge >1 and its associated hydrated ‘shell’ could potentially lead to switch-like functional outputs, perhaps including the regulation of catalytic output, or creation of unique binding sites for protein:protein or other biomolecular interactions.

Of mechanistic interest, we identified independent pHis-containing peptides from distinct protein kinases that aligned to the same region of the catalytic domain (the αC-β4 loop), which contains a phosphorylatable His-containing motif (HxN). MD simulations suggested extensive changes in the relative conformation of the αC helix with respect to the activation loop when 3-pHis was installed at this position in the cell cycle protein kinase PLK2. The structural importance of this region in protein kinases for modulating catalytic activity, and conservation of His throughout 67% of the ePK superfamily, hints that phosphorylation at the HxN motif could act as a more general regulatory mechanism. Studies are underway to evaluate this hypothesis.

The possibility of a new mode of competition between phosphate and the myriad of other PTMs that occur on Arg and Lys (most obviously methylation and acetylation) is note-worthy, and might have significant implications for understanding combinatorial cell signalling and epigenetics, particularly as histones are prone to phosphorylation on Arg and Lys as well as His^15^.

Validation of the huge number of novel phosphorylation sites reported here, and enhanced understanding of their physiological roles, will undoubtedly require the development of new tools and methodologies. The generation of generic and/or site-specific phospho-antibodies, alongside chemical genetic strategies for site-specific mimicry or genetic encoding of non-canonical phosphorylated residues, will be critical for mechanistic elucidation of function. Importantly, our discovery of the widespread nature of phosphorylation on multiple acid-labile PTMs, and the relative simplicity with which UPAX and MS approaches can be adopted for their analysis in complex mixtures, argues that the extent and biological relevance of non-canonical phosphorylation events for core and disease cell biology can be revealed quite rapidly.

A final experimental challenge will be the genomic annotation of the enzymes that catalyse or hydrolyse non-canonical phosphorylation, and biochemical and cellular analysis to help refine current understanding of eukaryotic signalling. The identification of protein domains that monitor or recognise non-canonical moieties (should they exist) are predicted to be employed for the assembly (or disassembly) of signalling networks. The diversity and prevalence of multiple non-canonical phosphorylation sites raises the question of how they contribute to global cell biology, and whether (as established for pTyr), the pX marks themselves, or the processes that phosphorylation controls, represent biomarkers, drug targets or antitargets in disease-associated signalling networks. In this context, it will be interesting to evaluate how non-canonical phosphorylation influences basic biological processes such as the cell cycle, ageing and apoptosis. Moreover, understanding the influence of drugs that impact signalling pathways, most notably the large number of small molecules that impact on protein phosphorylation, will also be of critical clinical importance.

## Methods

### Preparation of histidine phosphorylated myoglobin standard

Potassium phosphoramidate was synthesised from phosphoryl chloride and ammonia according to the procedure described by Wei and Matthews as described ^25, 26, 77^. In brief, phosphoryl chloride was reacted with ammonium hydroxide for 15 minutes on ice producing ammonium hydroxide phosphate, which was added to potassium hydroxide at 50 °C for 10 minutes. Potassium phosphoramidate (PPA) was precipitated with ethanol and collected by vacuum filtration. Equine myoglobin was phosphorylated by dissolution in 1 M aqueous PPA (150 nmol/mL) overnight at room temperature. Phosphorylation was evaluated by intact mass analysis of the resulting phosphorylated protein (5 μM in 20 mM ammonium acetate) by electrospray ionisation (ESI) direct infusion into a Synapt G2-S*i* mass spectrometer (Waters, UK).

### HeLa cell culture and siRNA knockdown of PHPT1

HeLa cells were maintained in DMEM (Sigma-Aldrich, Dulbecco’s Modified Eagle’s Medium-high glucose, 4500 mg/L glucose with sodium bicarbonate, without L-glutamine and sodium pyruvate) supplemented with 10% fetal bovine serum, penicillin (100 U/mL) and streptomycin (100 U/mL) at 37 °C, 5% CO_2_. To passage cells, cells were washed with warm PBS (Sigma-Aldrich, Phosphate Buffered Saline) prior to incubation with 1 mL trypsin (0.05% (v/v)) for 1 minute at 37 °C. Reaction was quenched with 1 mL supplemented DMEM. For siRNA knockdown T75 flasks at ^~^50% confluency were exchanged to antibiotic-free media (DMEM supplemented with 10% fetal bovine serum). siRNA for PHPT1 (SMARTpool: ON-TARGETplus), Lamin A/C (siGENOME control) and a non-targeting pool (ON-TARGETplus Non-targeting Pool) were purchased from Dharmacon. For each flask 1 nM siRNA (1.1 μL of a 20 μM stock prepared in RNAse free water (Thermo Fisher)) and 40 μL INTERFERin (Polyplus transfection) was prepared in 4 mL Opti-mem reduced serum media (Thermo Fisher). After 10 minutes incubation at room temperature siRNA mixtures were added to flasks. Cells were incubated for 48 hours at 37 °C, 5% CO_2_. For cell lysis, trypsinized cells were centrifuged at 220 *g* for 5 minutes, washed with PBS and lysed with 100 μL lysis buffer (8 M urea, 50 mM AmBic, 1 protease inhibitor tablet (cOmplete Mini EDTA free, Roche) per 10 mL). Lysate was sonicated at low amplitude for 3 × 10 seconds with a 1 minute gap. Protein concentration was determined by Bradford assay.

### SDS-PAGE and western blotting

Cell lysates from PHPT1 and Non-targeting (NT) siRNA experiments were diluted 1:2 with 2 X sample loading buffer (0.06 M Tris-HCl (pH 6.8), 10% (v/v) glycerol, 10% (w/v) SDS, 0.005% (v/v) bromophenol blue, 0.1 M DTT), and normalised for total protein loading. Samples were boiled for 5 minutes and loaded on a SDS-polyacrylamide gel electrophoresis gel. For western blotting, proteins were transferred to nitrocellulose, blocked and incubated with the appropriate primary antibody (PHPT1 (Santa Cruz Biotechnology, sc130229, 1:200 dilution); Lamin (Santa Cruz Biotechnology, sc6215, 1:200 dilution); GAPDH (manufacturer, 1:5000 dilution)) in blocking buffer overnight at 4°C. Signal was visualised with SuperSignal West Pico PLUS chemiluminescent substrate (Thermo Fisher) using X-Ray film.

### In-solution tryptic digestion

Phosphorylated protein standards, histidine-phosphorylated myoglobin (200 μg) and α- and β-casein (Sigma Aldrich, 100 μg of each), were dissolved in 25 mM ammonium bicarbonate (AmBic) to 2 μg/μL. Cell lysates were prepared in lysis buffer, as previously described. Proteins were reduced with 3 mM DTT (in 50 mM AmBic) for 20 minutes at 30 °C, and, after cooling, free Cys residues were alkylated with 14 mM iodoacetamide (in 50 mM AmBic) for 45 minutes at room temperature in the dark. The reaction was quenched by addition of DTT to final concentration of 7 mM. The urea concentration in cell lysates was reduced to 2 M by addition of 50 mM Ambic. Proteins were digested using 2% (w/w) sequencing grade modified trypsin (Promega) at 30 °C overnight.

### Titanium dioxide enrichment

TiO_2_ enrichment of α- and β- casein and histidine-phosphorylated myoglobin peptides (200 pmol) was performed using 200 μL spin tips (Protea Biosciences), with 3 sets of binding, wash and elution conditions as outlined in Supp. Table 2. Briefly, tips were prepared by addition of binding buffer (200 μL) and centrifuged at 2000 *g* for 1 minute. Peptides in 200 μL binding buffer were added to the tip, centrifuged, reloaded and centrifuged again. The resulting flow through (unbound material) was collected for analysis. Tips were washed with 200 μL each of binding and wash buffers, with each fractionation collected by centrifugation. Bound peptides were eluted by addition of 2 X 100 μL elution buffer. All fractions were dried by vacuum centrifugation and reconstituted in H_2_O:ACN (97:3) for LC-MS/MS analysis with the Bruker AmaZon instrument. The non-enriched peptides (‘start material’) were diluted to a concentration of 500 fmol/μL for LC-MS/MS analysis.

### Strong Anion Exchange (SAX) Chromatography

SAX was performed using a Dionex U3000 HPLC instrument equipped with a fraction collector. Peptides from phosphorylated protein standards (25 μg each α- and β- casein and histidine-phosphorylated myoglobin) or digested cell lysate (2 mg) were chromatographed using a PolySAX LP column (PolyLC; 4.6 mm inner diameter (i.d.) × 200 mm, 5 μm particle size, 300 Å) with a binary solvent system of solvent A (20 mM ammonium acetate, 10% ACN) and solvent B (300 mM triethylammonium phosphate, 10% ACN) at pH 6.0, pH 6.8 or pH 8.0. Solvent was delivered at 1 mL/min according to the following gradient: 5 minutes at 100%solvent A, then 43 minutes to 100% solvent B, and then 5 minutes at 100% solvent B before equilibration to start conditions. Fractions were collected every minute for 48 minutes, with every 3 pooled and the volume reduced by drying under vacuum to give 16 fractions in total.

### C18 StageTip desalting

StageTips were prepared with 3 discs of C18 material (manufacturer) in a 200 μL pipette tip. Tips were conditioned by sequential addition of 100 μL methanol, 100 μL H_2_O:ACN (50:50) and 100 μL H_2_O, with centrifugation for 2 minutes at 2000 rpm to pass the liquid through the tip each time. A portion of each SAX fraction (100 μL) was loaded onto the tip, centrifuged, then the flow through added to the tip and centrifuged again. The tip was washed with 100 μL H_2_O and peptides were then eluted by addition of 50 μL H_2_O:ACN (50:50). Eluents were dried to completion by vacuum centrifugation then resolubilised in H_2_O:ACN (97:3) prior to LC-MS/MS analysis.

### LC-MS/MS

#### AmaZon ETD

LC-MS/MS analysis of fractions obtained during pHis enrichment optimisation was performed using the AmaZon ETD ion trap (Bruker Daltonics, Bremen, Germany) arranged in-line with a nanoAcquity n-UHPLC system (Waters Ltd., Elstree, UK). Peptides were loaded from an autosampler onto a Symmetry C_18_ trapping column (5 μm packing material, 180 μm × 20 mm) (Waters Ltd., Elstree, UK) at a flow rate of 5 μL/min of solvent A (0.1% (v/v) formic acid in H_2_O), trapped for 3 min and then resolved on a nanoACQUITY C_18_ analytical column (1.8 μm packing material, 75 μm × 150 mm) (Waters Ltd., Elstree, UK) using a gradient of 97% A, 3% B (0.1% (v/v) formic acid in ACN) to 60% A, 40% B over 60 min at 300 nL/min. The column effluent was introduced into the AmaZon ETD ion trap mass spectrometer via a nano-ESI source with capillary voltage of 2.5 kV. Full scan ESI-MS spectra were acquired over 150-2000 *m*/*z*, with the three most abundant ions being selected for isolation and sequential activation by CID or ETD. A 1 min dynamic exclusion window was incorporated to avoid repeated isolation and fragmentation of the same precursor ion. CID was performed with helium as the target gas, with the MS/MS fragmentation amplitude set at 1.20 V, and ramped from 30 to 300% of the set value. For ETD, peptides were incubated with fluoranthene anions (ICC target 100000, max ETD reagent accumulation time 10 ms, ETD reaction time 100 ms).

#### Orbitrap Fusion

nLC-ESI-MS/MS analysis of cell lysate fractions was performed using an Orbitrap Fusion tribrid mass spectrometer (Thermo Scientific) attached to an UltiMate 3000 nano system (Dionex). Peptides were loaded onto the trapping column (Thermo Scientific, PepMap100, C18, 300 μm X 5 mm), using partial loop injection, for seven minutes at a flow rate of 9 μL/min with 2% ACN 0.1% (v/v) TFA and then resolved on an analytical column (Easy-Spray C18, 75 μm × 500 mm, 2 μm bead diameter) using a gradient of 96.2% A (0.1% formic acid in H_2_O) 3.8% B (0.1% formic acid in 80:20 ACN:H_2_O) to 50% B over 90 minutes at a flow rate of 300 nL/min. A full scan mass spectrum was acquired over *m*/*z* 350-2000 in the Orbitrap (120K resolution at *m*/*z* 200) and data-dependent MS/MS analysis performed using a top speed approach (cycle time of 3 s), with HCD (collision energy 32%, max injection time 35 ms) and neutral loss triggered (Δ98) EThcD (ETD reaction time 50 ms, max ETD reagent injection time 200 ms, supplemental activation energy 25%, max injection time 50 ms) for fragmentation. All product ions were detected in the ion trap (rapid mode).

### Proteomics data analysis

#### CompassXport/Mascot

MS output files from the Bruker AmaZon instrument were converted to .mgf files using CompassXport software (Bruker Daltonic) and an in-house script. The resulting .mgf files were searched using the Mascot search algorithm (version 2.6). Peptide and MS/MS mass tolerance: 0.6 Da; Database: Swissprot (2014.07.21); Taxonomy: mammalian; Missed cleavages: 2; Fixed modifications: carbamidomethyl (C); Variable modifications: oxidation (M), phosphorylation (ST), (Y) and (H). Data was manually inspected using DataAnalysis software (Bruker Daltonic) for the presence of phosphohistidine-containing peptides, and to extract peak area values for (phospho)peptides.

#### PEAKS 7.5

Label-free quantification of myoglobin peptides in SAX fractions (either with casein or spiked into U2OS lysate) was performed following analysis with the Orbitrap Fusion tribrid mass spectrometer. Raw files were searched using the PEAKS search engine against either an in-house database created by combining the Uniprot Bovine and Equine databases (2016.02.19) or the Uniprot human reviewed database (2015.12.02). Search parameters as follows: Parent mass error tolerance: 10 ppm; Fragment mass error tolerance: 0.6 Da; Enzyme: trypsin; Missed cleavages: 2; Fixed modifications: carbamidomethyl (C); Variable modifications: oxidation (M), phosphorylation (STY) and (HCDR); max variable PTM per peptide: 4

#### Proteome Discoverer 1.4

Raw files acquired on the Thermo Fusion mass spectrometer (HCD-neutral loss triggered EThcD method) were converted to .mzML using ProteoWizard’s msconvert tool in order to perform MS2-level deisotoping. The resulting files were then processed using Proteome Discoverer. For each raw file scans were split into those arising from HCD and EThcD events using a collision energy filter (HCD: min CE 0, max CE 34; EThcD: min CE 35, max CE 1000) to generate 2 separate .mgf files. These were searched using the Mascot search algorithm (version 2.6) against the Uniprot Human database (2015.12.02; 20,187 sequences), parameters were set as follows: MS1 tolerance of 10 ppm; MS2 mass tolerance of 0.6 Da; enzyme specificity set to trypsin with 2 missed cleavages allowed; fixed modification of carbamidomethylation (C) and variable modifications: oxidation (M), phosphorylation (ST), (Y) and (X) where X=H, K, R, D, or E. Instrument type: ESI-Quad-TOF for HCD files, CID+ETD for EThcD files. Phosphopeptides were additionally analysed using the ptmRS node, with the ‘treat all spectra as EThcD’ option selected for EThcD data. A peptide FDR filter of 5% was applied and all data files (both HCD and EThcD for each fraction) were exported to .csv for further processing. Peptides with rank >1 and those classified as ‘ambiguous’ or ‘rejected’ were removed. MS2 spectra corresponding to phosphopeptides were assessed for the presence of 3 neutral loss peaks (Δ80, Δ98 and Δ116) with a mass tolerance of 0.5 Da and an intensity cut-off of 5% compared to the base peak ion and given a ‘Triplet’ score of 0, 1, 2 or 3 depending on the number of neutral loss peaks identified.

### Molecular Dynamics Simulations

PDB structure 4I5P^78^ was used to model the PLK2 kinase (residues 71−354). The phosphate group was attached to the N1 or N3 atom of H139 residue through PyMOL. Due to the side chain packing of H139 in the crystal structure, a flipping of the imidazole ring is necessary before addition of the phosphate group to the N1 atom. The modified structures were then relaxed to Rosetta talaris2014 energy function and manually checked for steric clashes prior to molecular dynamics (MD) simulation^79^.

MD simulations were performed using GROMACS 5.0.4^80^. AMBER ff99SB-ildn force field^81^ was used with parameters for pHis adopted from previous quantum mechanics calculations^82^. Specifically, pHis-N1-H1D represents phosphohistidine at the N1 atom with single protonated phosphate group; pHis-N1-H2D represents phosphohistidine at the N1 atom with an unprotonated phosphate group; pHis-N3-H1E represents phosphohistidine at the N3 atom with single protonated phosphate group; pHis-N3-H2E represents phosphohistidine at the N3 atom with unprotonated protonated phosphate group. The protein was solvated with TIP3P water in a dodecahedron periodic box that was at least 1 nm larger than the protein on all sides. Sodium and chloride ions (0.1 M concentration) were then added to neutralize the net charge of the protein. Long-range electrostatics were calculated using Particle Mesh Ewald (PME) method with a neighbour-list cutoff of 9 Å. Energy minimization was carried out by coupling steepest descent and then conjugate gradient algorithms until F_max_ was <100 kcal/mol. Temperature equilibrium was achieved using velocity rescaling with a stochastic term for 200 ps at 310 K^83^. NPT ensemble was done for 200 ps at 1.0 bar using berendsen exponential relaxation pressure coupling. The unrestrained MD were run for 500 ns using a time step of 2 fs after NPT equilibrium. Root mean square deviation (RMSD) of each simulation was checked for stability before additional analysis. Built-in utilities of gromacs such as, g_mindist and g_rmsf, were used to process the trajectory data. Free energy landscape plot was generated using MSMExplorer^84^. All protein visualization was done using PyMOL.

## Acknowledgements

This work was supported by funding from the Biotechnology and Biological Sciences Research Council (BBSRC) to C.E. (BB/H007113/1, BB/M012557/1), North West Cancer Research (CR1037 and CR1088) and a BBSRC DTP PhD studentship to G.H.

